# Stratified Active Learning for Spatiotemporal Generalisation in Bioacoustic Monitoring

**DOI:** 10.1101/2025.09.01.673472

**Authors:** Ben McEwen, Corentin Bernard, Dan Stowell

**Affiliations:** Tilburg University, Netherlands; Toulon University, France; Naturalis Biodiversity Center, Netherlands

**Keywords:** Active Learning, Audio Classification, Bioacoustics, Stratification

## Abstract

Active learning optimises machine learning model training through the data-efficient selection of informative samples for annotation and training. In the context of biodiversity monitoring using passive acoustic monitoring, active learning offers a promising strategy to reduce the fundamental annotation bottleneck and improve global training efficiency. However, the generalisability of model performance across ecologically relevant strata (e.g. sites, season etc) is often overlooked. As passive acoustic monitoring is extended to larger scales and finer resolutions, inter-strata spatiotemporal variability also increases. We introduce and investigate the concept of *stratified active learning* to achieve reliable and generalisable model performance across deployment conditions. We compare between implicit cluster-based diversification methods and explicit stratification, demonstrating that cross-strata generalisation is a function of stratum divergence, not sampling balance. Additionally, mutual information as well as exclusion analysis show that spatiotemporal context can explain a substantial proportion of species label variance and inform sampling decisions.

## 1. Introduction

Passive acoustic monitoring (PAM) provides biodiversity insights at previously infeasible scales and resolutions [1, 2]. The affordability of acoustic recording devices (ARDs) combined with recent improvements such as network-connected data streaming vastly improve the scalability of monitoring but also the quantity of raw data collected [3]. Recent gains in computational bioacoustics [4] are fundamentally data-driven, with the quality of inference dependent on the quality and quantity of labelled data. However, as the quantity of raw acoustic data increases exponentially the availability of data labels remains a severe bottleneck, due to the requirement of skilled annotators.

Further, as biodiversity monitoring expands to larger spatial and temporal scales, the ecological and acoustic heterogeneity across recording sites increases. Monitoring stations may be deployed across diverse habitats and regions where differences in species composition and background noise, such as wind, rain, insect activity and anthropogenic sounds, introduce significant spatial variation. Different habitats also affect sound attenuation [5, 6] and model detectability [7]. The detectability of certain species also changes with seasonal and environmental factors [8] and also variation due to regional dialects [9]. There is also potential variability between recording devices [10] which makes generalisable performance across locations more challenging. Temporal variability further arises from daily vocalisation patterns (e.g. dawn chorus), seasonal dynamics driven by phenology and migration, and long-term changes dues to microphone degradation or shifting climate and habitat conditions. Models for species recognition are expected to produce reliable predictions across this full range of conditions, requiring that training data is representative of the underlying variability across deployment conditions.

Active learning is the machine learning technique of dynamically selecting informative samples for annotation and model training. By allocating annotation resources to high-priority samples, active learning aims to achieve higher model performance with fewer annotated samples (training efficiency). Active learning, based on uncertainty sampling, has been demonstrated to improve global training efficiency (peak performance per sample) [11] as well as per species training efficiency [12, 13].

Sample diversification is an important aspect of active learning. To maximise the information gain, samples must have high entropy/uncertainty (i.e. not already predictable) *and* diversity. Diversification minimises redundant information. For practical purposes, active learning is typically applied over batches. Larger batch sizes reduce adaptivity and increase the risk of including redundant samples [14, 15]. Methods such as batched active learning, where the batch size refers to the number of samples selected for annotation per iteration, implicitly promote sample diversity across batches by iteratively updating the modeled probability distribution given a reduction in entropy. However, this does not account for intra-batch diversity and so various hybrid approaches exist to improve intra-batch diversity [16]. Diversification alone is not effective as samples should be both informative (high-entropy) and non-redundant (not currently represented in the batch) [17]. Hybrid sampling approaches that attempt to select both informative but diverse samples have demonstrated improved performance compared to pure diversification methods [13].

Clustering is generally applied to improve the diversity of sample selection by sub-sampling clusters within an embedding space (i.e. based on acoustic features). High-quality embeddings capture relevant variability without explicitly using spatial or temporal information. Stratified sampling, by explicitly selecting samples from defined strata such as sites, seasons, or years, is a common way to ensure training data captures ecological variability across space and time (Figure 1). Stratification can be considered a form of explicit diversification, in contrast to traditional active learning diversification methods which implicitly account for variability.

**Figure 1:**
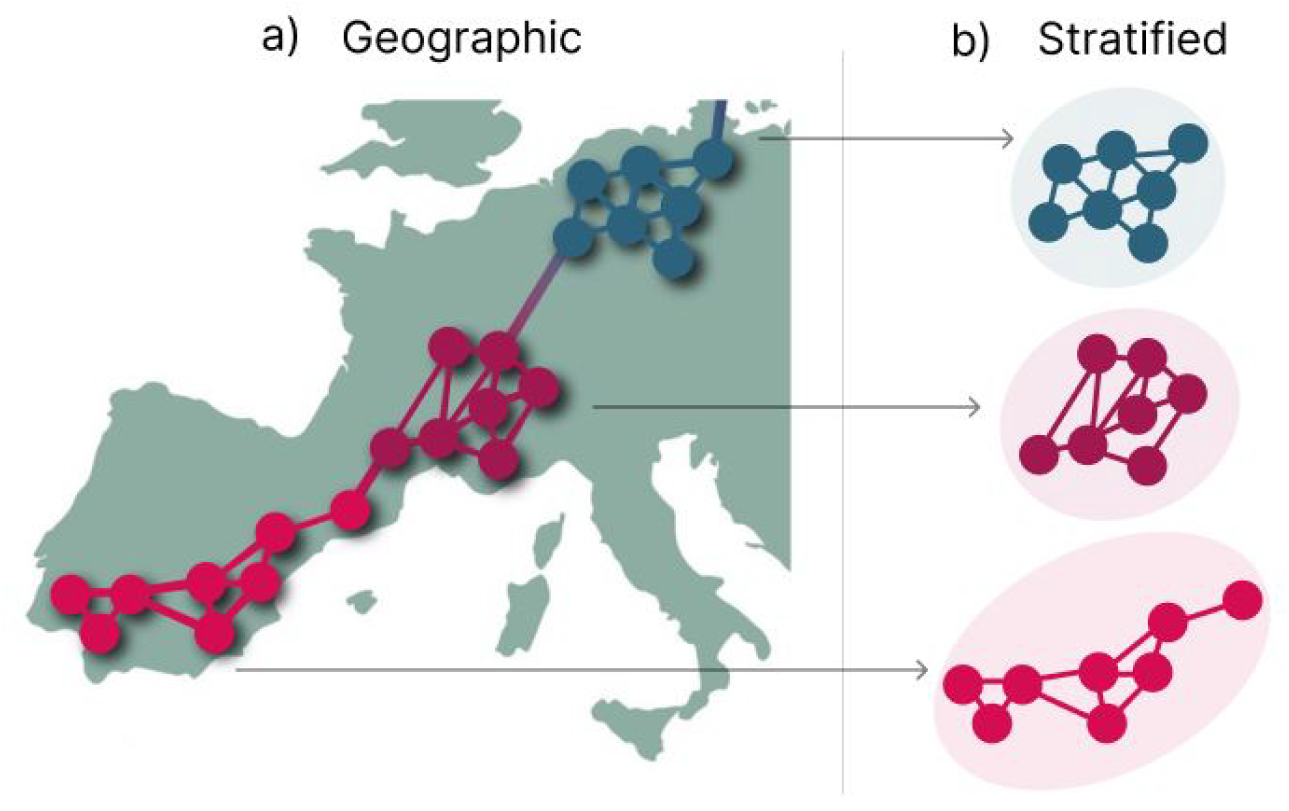
Demonstration of stratified sampling (b) based on geographic location (a).

## 2. Contributions

This paper makes the following contributions:

- We demonstrate that model performance, using traditional transfer learning and active learning, does not generalise across monitoring sites.
- We develop a framework for evaluating model and sampling performance across spatial and temporal strata.
- We empiricially compare implicit and explicit active learning diversification methods on relevant multi-site long-term datasets and propose the method of *stratified active learning*.
- We analyse stratum-level divergence, investigating methods to identify mutual information between strata.

In the following, we formalise our machine learning framework, before proposing the modifications of *stratified active learning*. We empirically evaluate this using public bioacoustic datasets of birds and anurans. Datasets are publicly available via their respective authors. All code notebooks are open-source and publiclyd [available](https://github.com/BenMcEwen1/Stratified-Sampling). Here, we focus on the application of acoustic monitoring but the contributions of this work are relevant across monitoring modalities.

## 3. Method

To summarise the method, we compare the performance evenness across strata using implicit cluster-based and explicit stratification-based diversification within an active learning framework. Performance is assessed using the aggregated F1 score per stratum with evenness quantified using the Pielou Evenness Index [18] and coefficient of variation, providing a framework for comparing model generalisability across ecological gradients. Additionally, we analyse stratum-level divergence using the Jensen-Shannon divergence and quantify the mutual information between strata with both label and predictive distributions.

### 3.1. Model

As input we use a set of embeddings pre-generated by a neural network (in our case, BirdNet [19] penultimate layer) 𝒳 = {*x*_1_, *x*_2_, …, *x*_*N*_ } where *x*_*n*_ ∈ ℝ^*N ×*1024^ is the embedding of a single acoustic obseration and 1024 is the feature dimension. A simple classifier model *f*_*θ*_(*x*_*n*_) is defined as single linear layer *f*_*θ*_(*x*_*n*_) : ℝ^1024^ → ℝ^|*c*|^ with an output dimension corresponding to the number of classes *c* using a standard transfer learning configuration. The efficacy of transfer learning, especially in the context of biodiversity monitoring application, has been demonstrated in various studies [20, 21, 12, 13]. The model generates a multi-label prediction, hence sigmoid activation is applied to the model logits *ŷ*_*n*_ = *σ*(*f*_*θ*_(*x*_*n*_)). Only the classification head is fine-tuned. The model and training hyperparameters are fixed across this study to isolate sampling effects, consistent with previous works [12, 13].

### 3.2. Sampling for Active Learning

Given samples across large spatial and temporal scales we want to compare if hybrid uncertainty sampling implicitly provides generalisable performance across strata or if explicit stratification improves performance. We use a standard uncertainty sampling approach, where the samples with the highest uncertainty are selected for annotation. Binary entropy is selected for uncertainty quantification and is a standard method for multi-label data hence this method is fixed across all sampling and diversification methods:

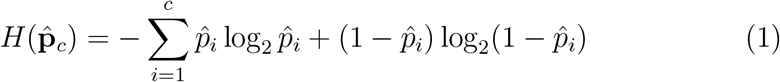

Per sample entropy is aggregated using the maximum across classes 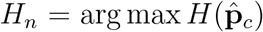 for each sample. Per-stratum binary entropy (without diversification) is reported as entropy sampling and is the baseline sampling method for comparison.

### 3.3. Diversification

We evaluate three sampling methods: 1) binary entropy without diversification (equation 1), 2) cluster-entropy diversification and 3) stratification-entropy diversification.

#### Cluster-Entropy Diversification

Clustering (Cluster-Entropy) using pre-generated BirdNet embeddings provide a content-based representation, allowing similar samples to be grouped. By sampling from clusters based on uncertainty, active learning can select informative examples while ensuring diversity across the embedding space, thereby reducing redundancy in the labeled set. Unlike stratification, where samples are explicitly categorised by site and date (year or month), cluster-based diversification is not provided with spatiotemporal information but embeddings may implicitly represent spatial and temporal information due to correlation with acoustic variability (e.g. species composition and environmental noise).

Based on a similar approach cluster-margin [15], cluster-entropy is a hybrid uncertainty sampling and diversification approach. A subset of sample embeddings (20%) are used to compute Hierarchical Agglomerative Clusters (HAC) using Ward linkage criteria for a precomputed set of clusters 𝒞_*cluster*_ = {*c*_1_, *c*_2_, …, *c*_*k*_} (depending on the desired sampling batch size *k* = 20) where each cluster *c* contains sample embeddings. All training embeddings are assigned to cluster means by Euclidean distance. Binary entropy is then computed for all clustered embeddings and the highest uncertainty samples from each cluster are selected:

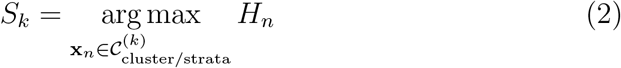

#### Stratified-Entropy Diversification

Samples are categorised by *t* strata varying by the number of sites (spatial) or time periods (temporal). Binary entropy is used to compute the uncertainty of samples 𝒞_*strata*_ = {*c*_1_, *c*_2_, …, *c*_*t*_ }. Similarly, the samples with the highest binary entropy uncertainty per stratum are selected (equation 2).

For stratified-entropy spatial (sites) and temporal (yearly, monthly) stratification are compared separately. The number of strata depends on the number of locations or time-periods present in the dataset. Therefore we apply posthoc random subsampling so that the sampling batch size remains consistent between compared approaches.

##### Hyperparameters

An iterative batched active learning process of up to 500 cumulative samples (25 epochs) were computed with 20 samples per batch for all sampling approaches. A mini-batch size of 20 was used with an Adam optimiser and learning rate scheduler (ReduceLROnPlateau) starting at 0.01, limited at a minimum of 0.0001.

### 3.4. Datasets

Datasets were selected which cover varied spatial and temporal scales as well as having necessary metadata regarding recording sites and dates. **AnuraSet** [22] is a multilabel bioacoustic dataset from South America containing vocalisation of 42 anura species. The dataset spans four recording sites and three years (2019 - 2021). This dataset reflect a multi-site national monitoring application. The World Annotated Bird Acoustic Dataset **(WABAD - Europe)** [23] is the most comprehensive multi-label bioacoustic dataset currently available. Recordings span 26 sites across Europe and seven years (2016 - 2023) reflecting a large-scale transnational monitoring study.

#### Challenges and Limitations

These datasets reflect samples collected from complex and real-world passive acoustic monitoring applications. These are challenging datasets given class imbalance and uneven sampling across strata (sites and time periods). These challenges reflect the real-world challenges of passive acoustic monitoring at scale. There are limitations when making comparison to raw field recordings. Sampling bias and over-representation of positive samples inflates the performance of non-diversification based sampling. These dataset do not reflect the true sparsity of positive samples. Evaluation of sampling strategies given a dataset with existing sampling biases is a point of caution. However, relative performance differences are still applicable.

### 3.5. Evaluation

The F1 (macro averaged by class) is the primary model performance metric computed on the test set. Macro (non-weighted average) metric was selected because it more accurately represent performance for underrepresented species. Performance is calculated per stratum. A fixed threshold of 0.5 is used for all F1 evaluation.

An important and often overlooked aspect of evaluation is the generalisability of model performance, especially across spatial and temporal scales with significant variability. Intra and inter-strata evenness of performance metrics are evaluated using the Pielou Evenness Index *J ′* [18] (also called the Shannon Equitability Index), which computes the sum of per stratum entropy over the maximum entropy:

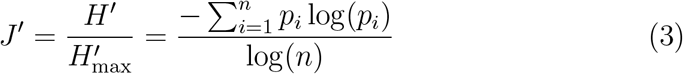

The Pielou Index ranges from 0 to 1 with a higher value indicating better evenness. Additionally, the coefficient of variation (CV), equation (4), is reported which measures the relative dispersion of performance across strata where lower values indicate better evenness:

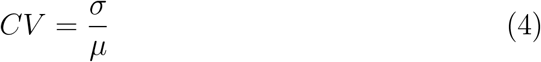

Here we are defining performance evenness across strata as a proxy for generalisability. True generalisability also implies robustness beyond observed strata. Evenness can also be considered as an indicator of fairness across sub-populations.

### 3.6. Divergence and Mutual Information

Jensen-Shannon (JS) Divergence is used to measure distributional differences between strata by computing the pair-wise JS-divergence. The JS-divergence of the true label distributions and predicted class distributions (categorical distributions) are computed, estimating the true distributional differences between strata and the differences captured by the model respectively.

The mutual information quantifies the amount of information that knowing one variable (e.g. stratum S) reveals about the other (e.g. prediction or label Y). The normalised mutual information is used to compare the proportion of uncertainty explained by strata information:

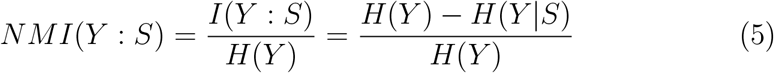

In this case Y is a categorical distribution across classes, either the true distribution (label) or predictive distribution. These divergence measures are used to explore potential explanations for the performance evaluation outcomes.

### 3.7. Exclusion Analysis

Exclusion analysis provides a direct evaluation of model generalisability across strata. By withholding samples from all but one stratum we evaluate the generalisability of the model across all strata and assess how well the model transfers to unseen but ecologically relevant conditions. Here the relative F1 (macro averaged) performance is reported for each stratum which is the relative difference between the first and last (25th) epoch. This analysis is paired with JS-divergence measurements between strata to measure the distribution divergence (predicted performance divergence) across strata.

## 4. Results

### 4.1. Spatial Stratification

We first evaluate spatial stratification using the AnuraSet dataset evaluated across four sites (INCT20955, INCT4, INCT41, INCT17). Using binary entropy sampling without any diversification as a baseline, a Pielou Index of 0.968 and coeficient of variation of 0.290 was achieved.

Performance varied substantially across sites and correlated with the number of samples selected (Figure 2). The mean performance across strata is denoted by the blue dashed line. Note that in this case, per-stratum F1-macro (test) is only evaluative, no stratification of training samples is applied.

**Figure 2:**
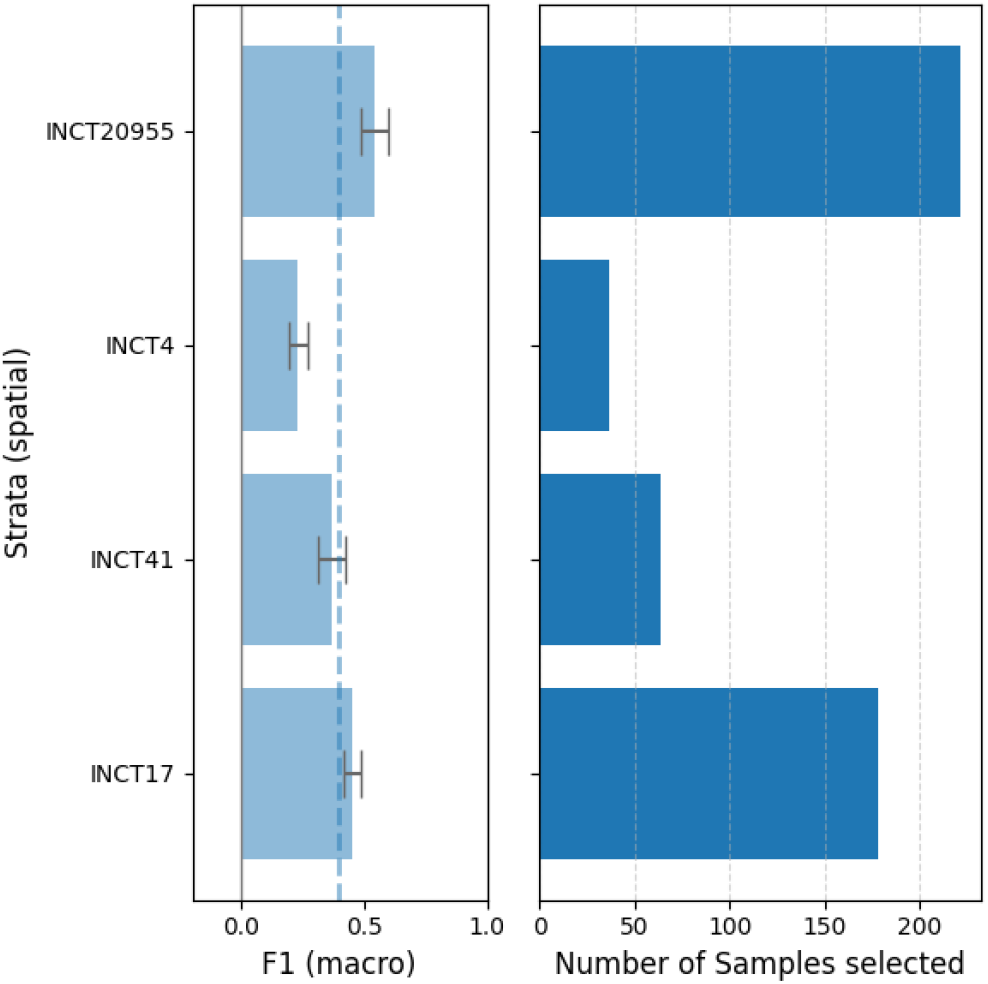
Comparison of model performance using AnuraSet with spatial stratification (sites) and entropy sampling without diversification.

#### Diversification comparison

When diversification was applied, both cluster-entropy and stratified-entropy improved performance evenness. Stratified-entropy sampling achieved a Pielou Index of 0.997 and coefficient of variation of 0.083. Cluster-based sampling achieved a Pielou Index of 0.994 and a coefficient of variation of 0.130 (Figure 3).

**Figure 3:**
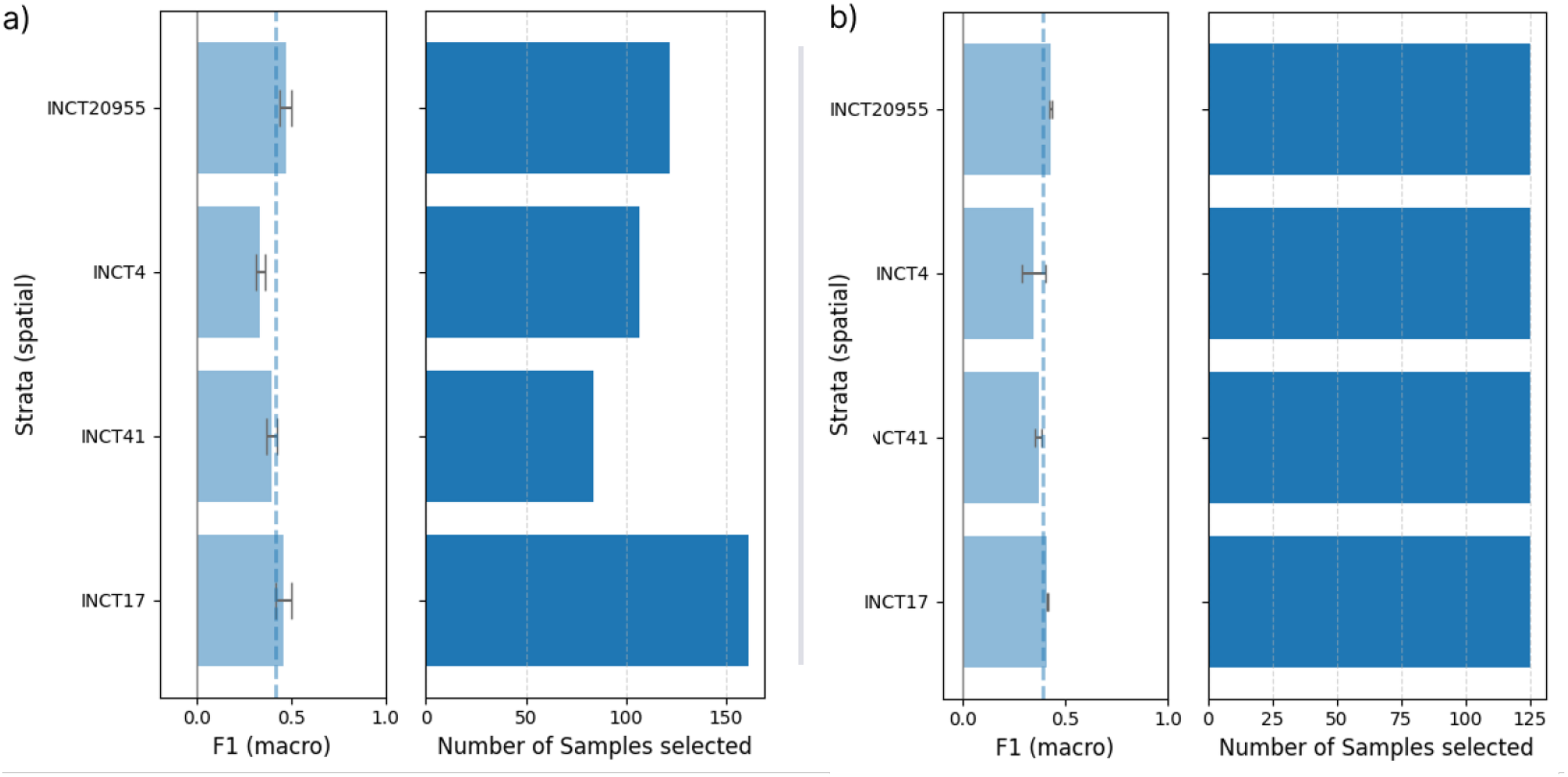
Comparison of cluster-entropy and stratified-entropy diversification using AnuraSet with spatial (site) stratification.

#### Exclusion Analysis

The exclusion analysis, where the model is evaluated across all site given only training data selected from one, showed lower performance on sites with greater divergence (Figure 4a). The corresponding pairwise JS-divegence between sites (Figure 4b) confirms a relationship between predictive divergence and cross-site generalisation (lighter colours denote higher dissimilarity).

**Figure 4:**
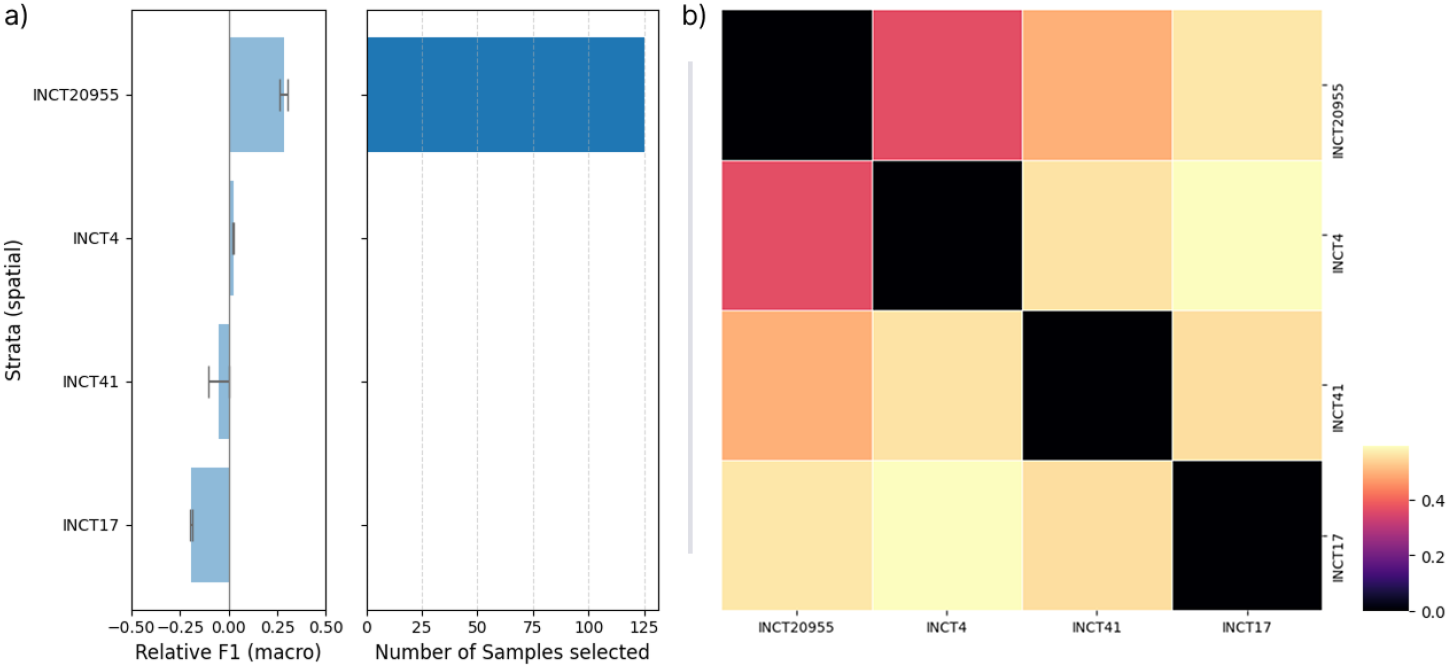
Exclusion analysis with sampling from a single site (INCT20955) showing a) per stratum performance and b) JS-divergence of predictive distribution between sites.

#### Mutual information

Mutual information quantifies the amount of information provided about the label given knowledge about the stratum (e.g. site, year, month). High mutual information indicates that the strata is informative of species composition (label distribution) or predicted labels (predictive distribution). For AnuraSet (spatial) the mutual information between the strata and species true label distribution is 40.3% of the total entropy. After training, the mutual information of the predictive distribution with respect to the strata reaches 34.9%, approaching the upper-bound of 40.3%. This suggests that spatial information explains a substantial portion of label variability and that species composition varies meaningfully across sites discussed further in (Section 5.1).

### 4.2. Temporal Stratification

Temporal stratification was evaluated on the same dataset (AnuraSet) across three years of data (2019 - 2021). Using entropy and cluster-entropy sampling it is observed that 2021 is undersampled (yellow region) yet the performance is even across all years (Figure 5).

**Figure 5:**
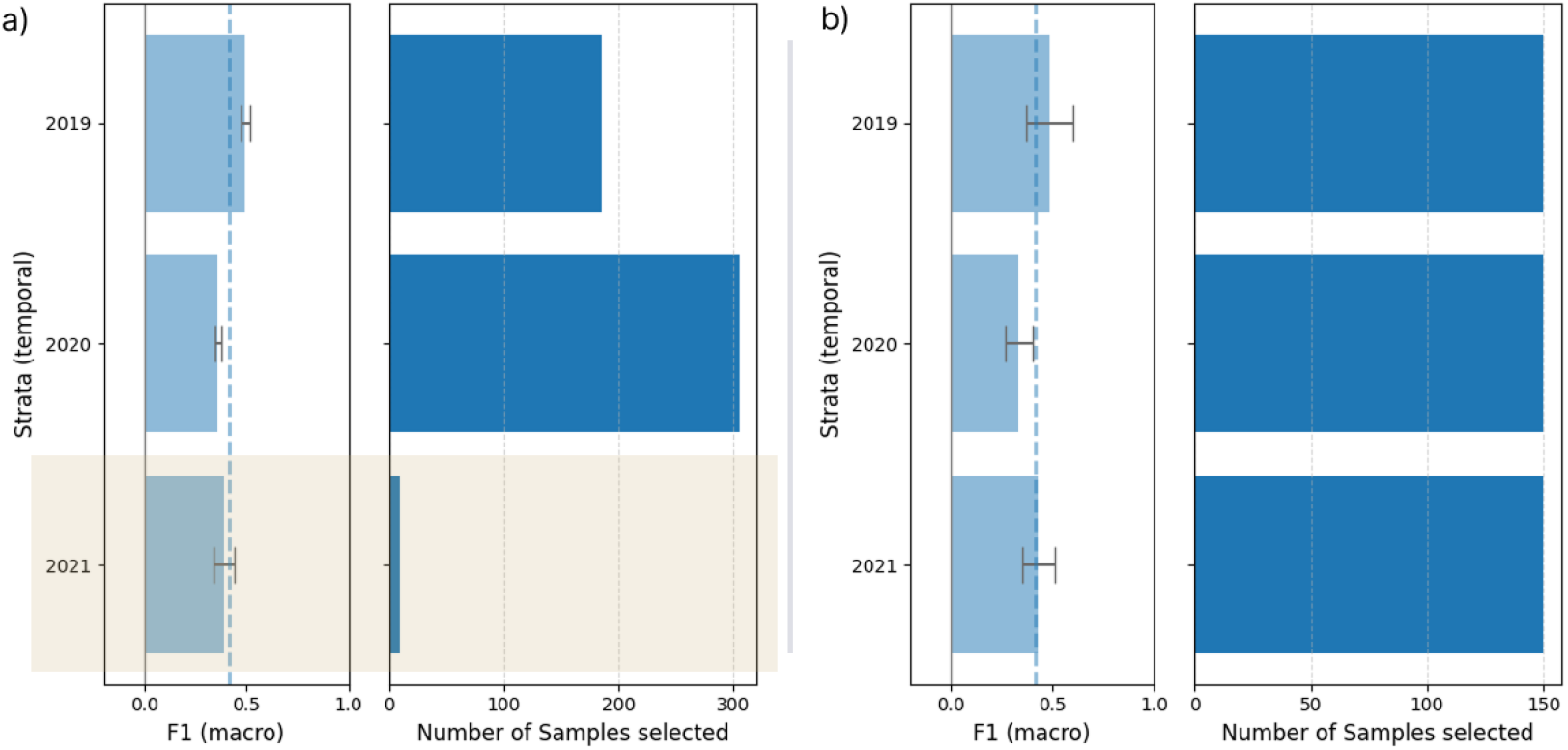
Comparison of a) cluster-entropy and b) stratified-entropy diversification using AnuraSet with temporal (year) stratification.

However, JS-divergence analysis revealed higher divergence for 2021 compared to other years. The mutual information between years and species labels is 6.18% (5.22% excluding 2021), with a predictive mutual information of 3.85% (3.36% excluding 2021). This discrepancy is discussed in Section 5.1

At a finer temporal resolution (monthly), consecutive months displayed low JS-divergence, and the mutual information between month and labels increased to 24.6% (predictive 15.3%), also suggesting that more relevant information has been captured at higher resolutions.

#### Sampling Resolution

When redefining custom date ranges based on the observed divergence patterns using custom date ranges {A, B, C} corresponding to {201909 to 201911, 201912 to 202004, 202010 to 202101} based on divergence from figure 6. This results in an improved evenness with a Pielou Index of 0.996 and coefficient of variation of 0.091. Model performance and divergence across custom date ranges is shown in Figure 7. Additionally, the mutual information between predictive distributions also increases from 3.85% to 16.0%. These results indicate that this custom resolution more effectively captures ecologically relevant variation, discussed further in Section 5.1.

**Figure 6:**
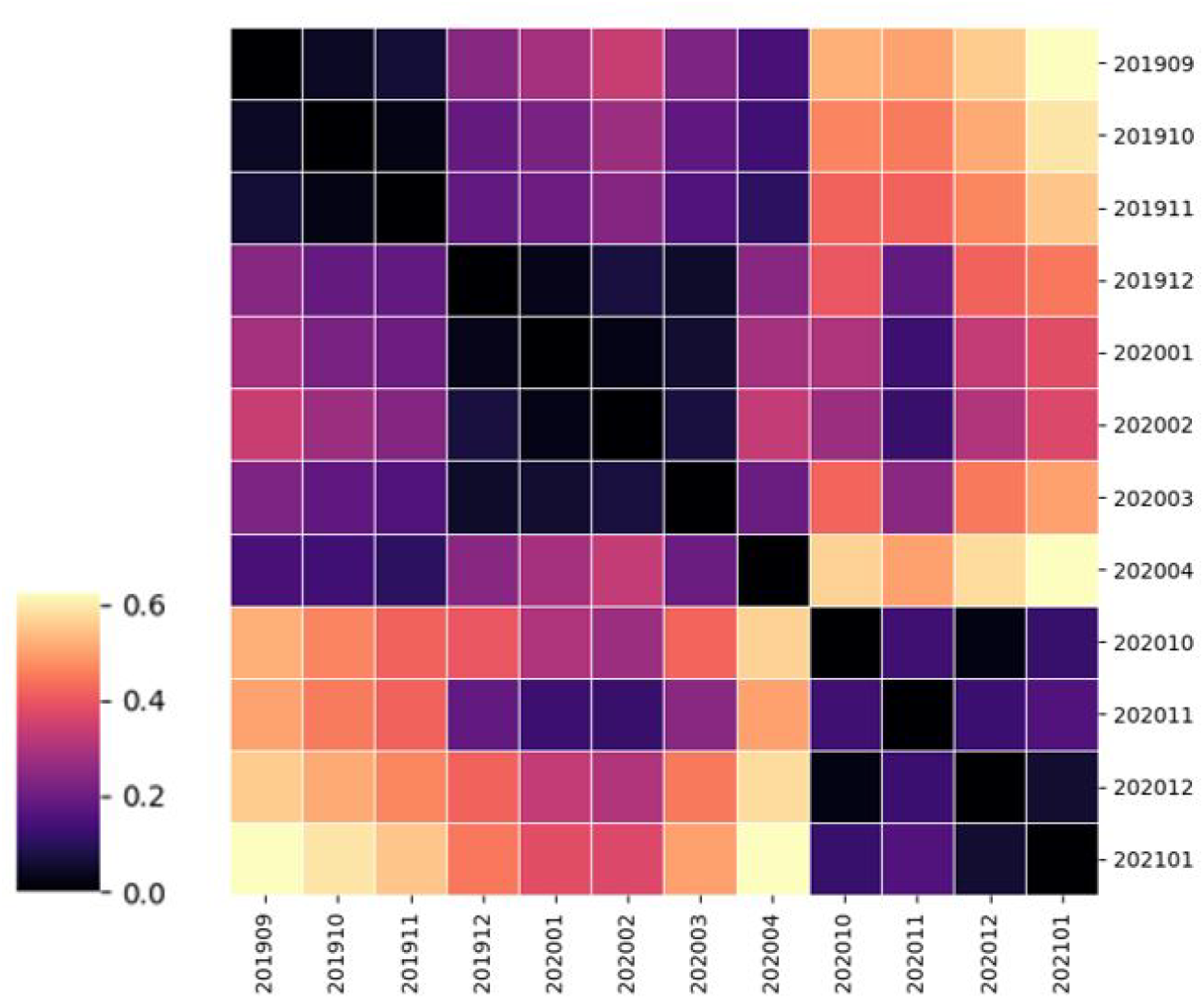
JS-divergence of per-month label distributions (AnuraSet)

**Figure 7:**
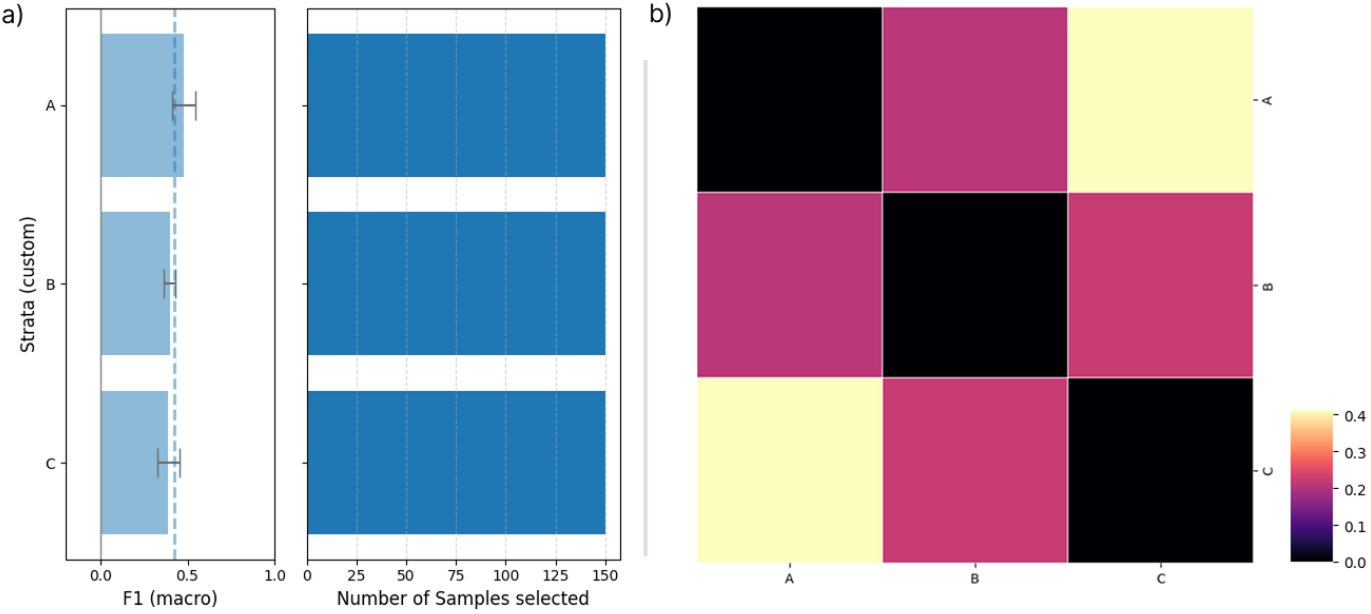
Comparison of a) per stratum performance and b) JS-divergence of predictive distribution between date ranges.

### 4.3. Scalability

Extending this evaluation to WABAD-Europe, which spans across 26 locations and 7 years (2016-2023) reveals high variability in model performance across spatially and temporally diverse recordings.

#### Spatial Stratification

Stratified sampling results in comparable performance evenness with a Pielou Index of 0.980 and coefficient of variation of 0.364 compared to cluster-entropy of 0.984 and 0.326 respectively. Both methods improve upon entropy sampling which achieved a Pielou Evenness of 0.970 and coefficient of variation of 0.435. Uneven sampling for stratified-entropy (Figure 8b), is due to posthoc subsampling required for balanced batchsizes.

**Figure 8:**
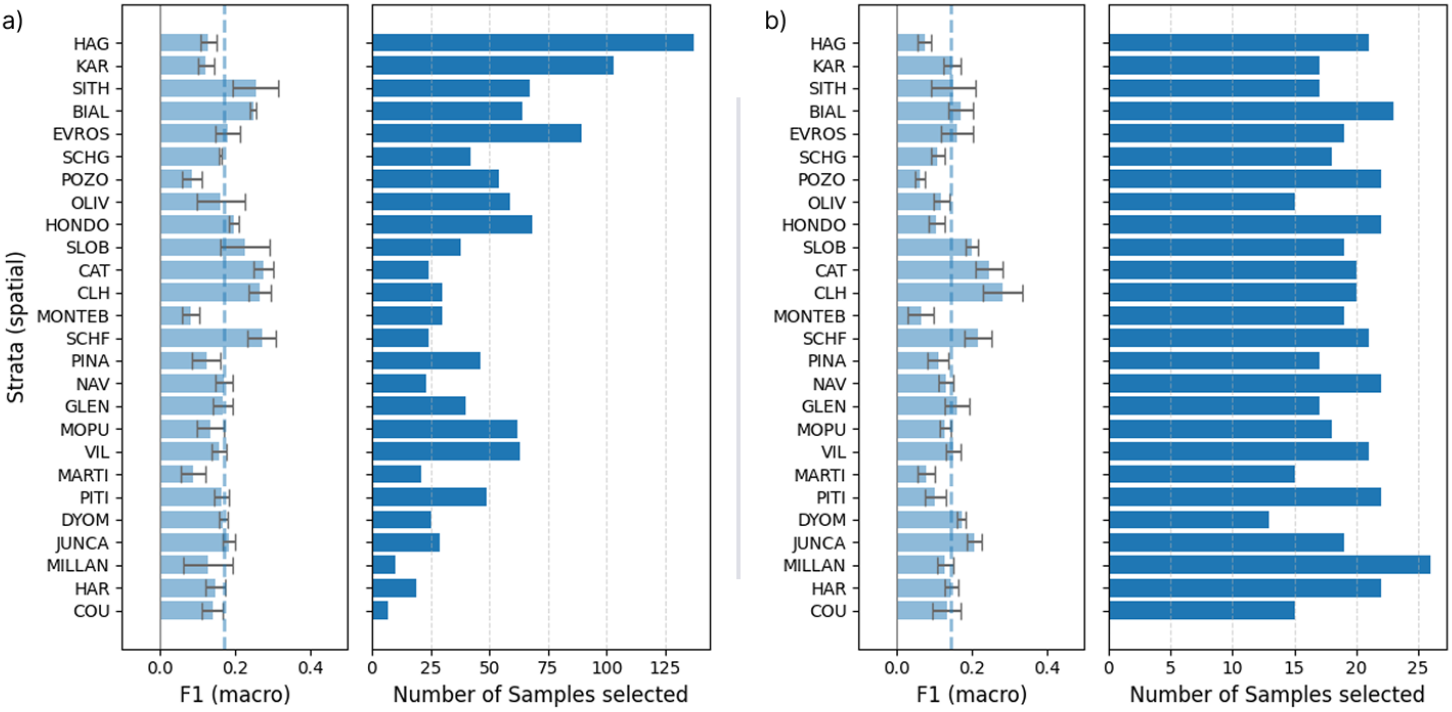
Comparable performance between a) cluster-entropy diversification and b) stratification.

The pairwise JS-divergence is shown across WABAD european sites (Figure 9). A large amount of variablity in the divergence across sites is observed. Comparing temporal stratification on WABAD-Europe highlights the same issue of imbalanced and inconsistent representation. Temporal stratification fails to improve performance evenness. This is again due to inconsistent month-level coverage and more specific to the WABAD dataset, the aggregation of recordings from many independent studies.

**Figure 9:**
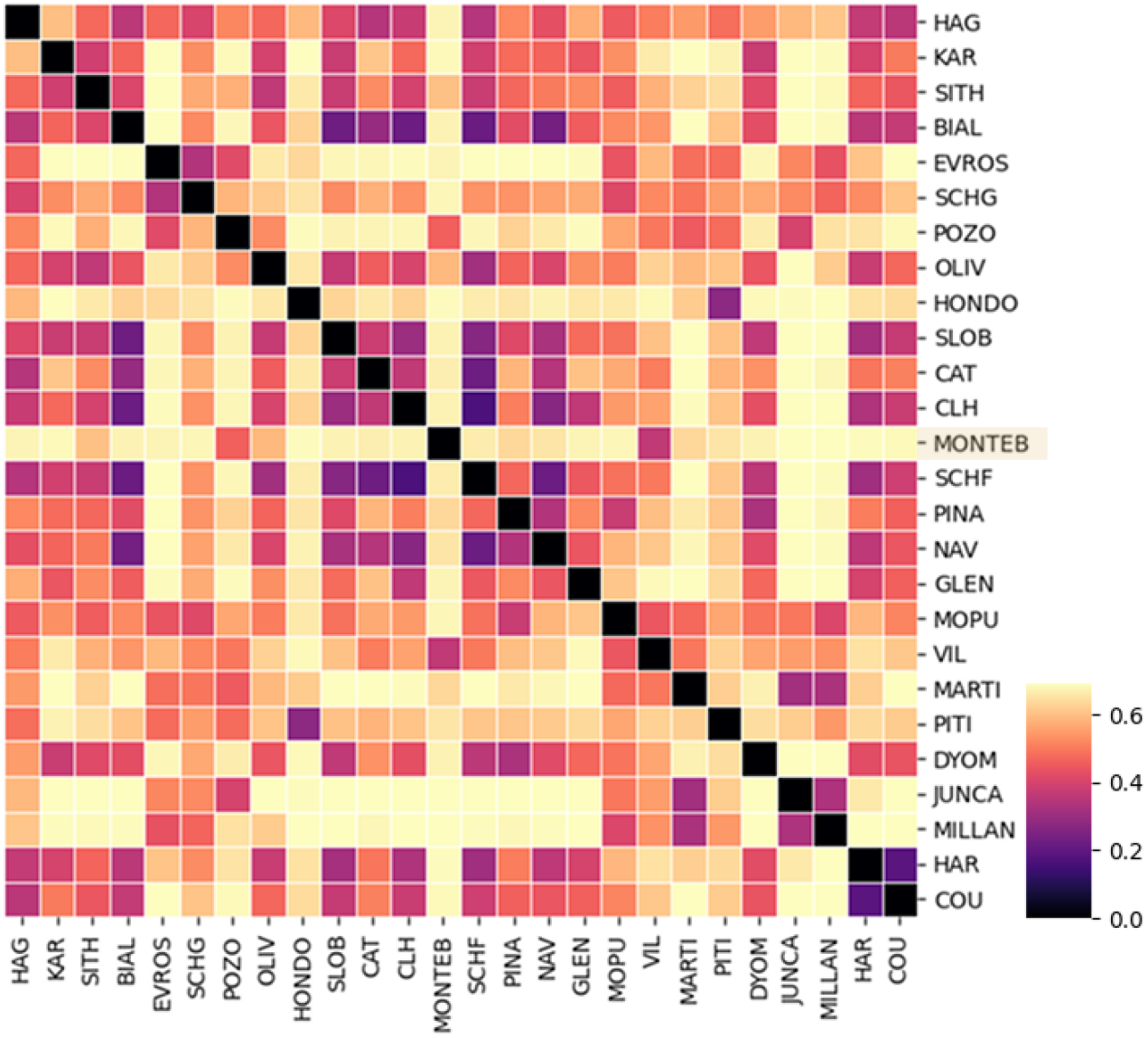
JS-divergence of per-site label distributions (WABAD-Europe)

**Figure 10:**
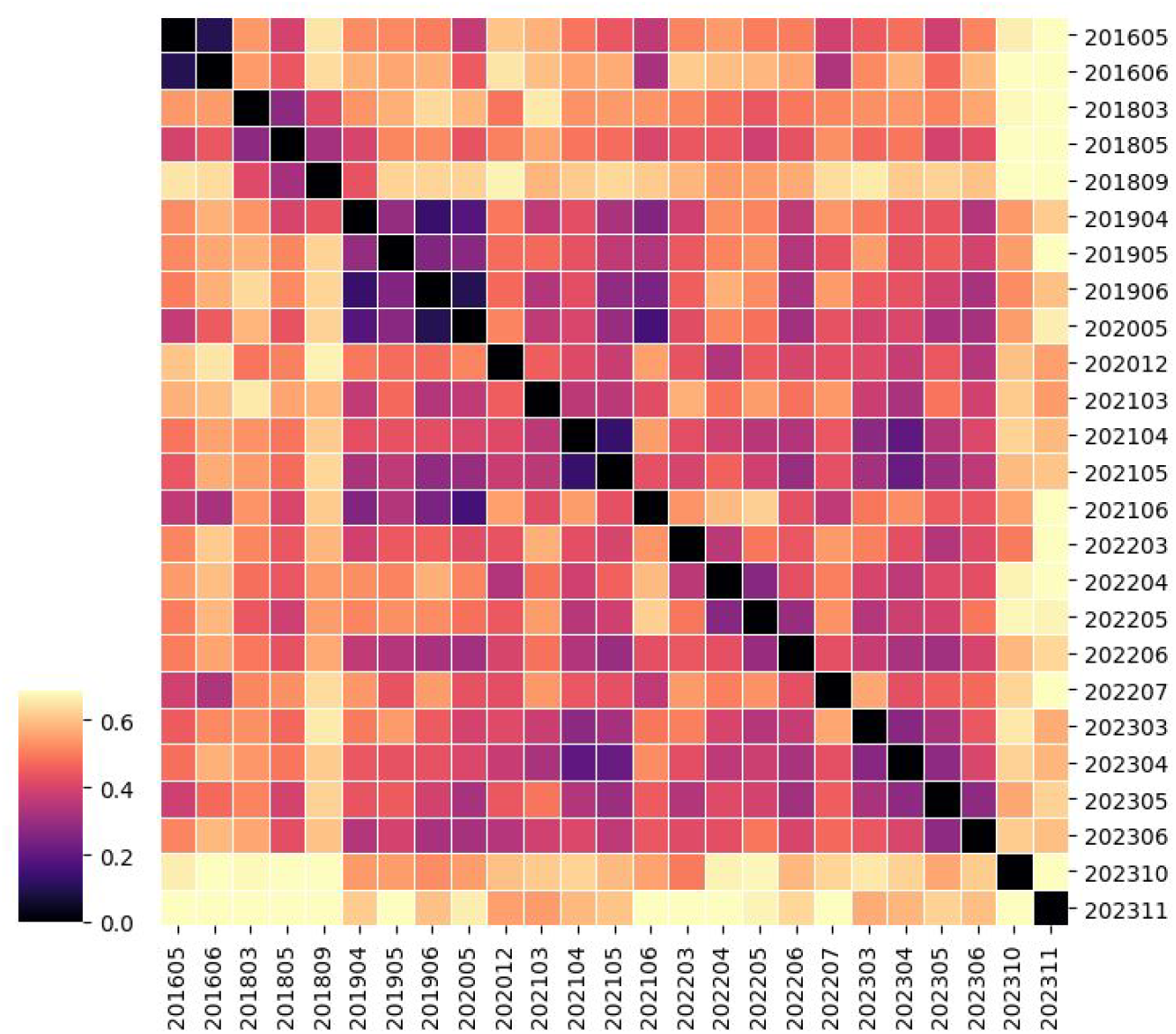
JS-divergence of per-month label distributions (WABAD-Europe)

## 5. Discussion

### 5.1. Spatial and Temporal Stratification

Balanced sampling across ecologically relevant spatial strata improved model performance evenness, confirming that sample diversity can enhance generalisability. However, uniform sampling alone is insufficient as demonstrated in Section 4.2 where even sampling across unrepresentative stratum reduced overall performance and evenness. *Stratification with balanced sampling introduces redundant samples, resulting in lower performance evenness*. This result would suggest that 2021 contains little relevant variation compared to other years. However, the JS-divergence analysis of the temporally stratified label distributions contradict this: 2021 has the highest divergence compared to other years. Inconsistencies in generalisation and divergence between years are due to **mismatched temporal coverage**, with 2021 only representing January. This leads to an artificial alignment with 2020 data, improving generalisation, while also exaggerating divergence, when a full-year distribution and partial-year distribution are compared. The JS-divergence Figure 6 demonstrates high-resolution (monthly) divergence.

Additionally, while it is true that diverse and representative data is a necessary component of model generalisability, a fundamental assumption when applying active learning is that samples are not uniformly informative. Therefore, model generalisability depends not just on sampling uniformly across strata, but also how informative those samples are. Ideally, samples are both representative and informative. Existing uncertainty quantification methods such as binary entropy provide a means to predict sample information gain and therefore inform sampling decisions.

### 5.2. Divergence-Informed Sampling

*Stratum Selection*. For spatial stratification, high mutual information between site and species labels show that location-based sampling captures relevant ecological information. This suggests that explicitly encoding spatial information into model embeddings (i.e. combining encoded inputs and encoded spatial information) could further improve performance.

Likewise, the monthly-level stratification and divergence-informed stratification produced high mutual information also implying temporal sampling can capture relevant variation. In contrast, a lower mutual information as well as model performance and evenness was observed using yearly stratification. This suggests that applying stratification across irrelevant or erronous axes and resolutions can harm model generalisability.

Ideally, stratification axes and resolution should align with underlying ecologically relevant information and cycles. In the case of these datasets it is likely that these results also reflect sampling and data collection decisions, not necessarily the underlying ecological information. This can be observed in Figure 6 where low divergence is observed across consecutive months but high-divergence is observed between the same month/season aross multiple years.

### 5.3. Scalability and Generalisation

Scaling to the larger WABAD-Europe dataset show that both cluster and stratification-based diversification yield only modest performance evenness gains and overall generalisability remains low. This is likely compounded by data imbalance and the datasets construction from many independent studies which introduce inconsistencies in spatial and temporal coverage. The WABAD-Europe dataset aggregates recordings from multiple independent projects. Such heterogenity likely obscures meaningful spatiotemporal structure.

### 5.4. Broader Implication and Limitations

The results presented highlight that stratified active learning can improve the evenness of model performance across ecological gradients, but its **success depends critically on how well the chosen stratification axes and resolution captures relevant variation**. Stratification by ecologically meaningful axes such as site or month enhances generalisability, while coarse or uninformative divisions, such as arbitrary yearly bins, can introduce redundancy and even degrade performance. **Measures such as JS-divergence and mutual information provide useful metrics by which to inform sampling decisions**. Interpretation must be made with caution: divergence can also reflect sampling biases, noise conditions, or inconsistent annotation effort rather than relevant information.

Notably, explicit stratification across a single axis (e.g. site or month) can out-perform cluster-based diversification methods. This is an interesting finding due to the embedding-space capturing richer semantic information related the sound source and potentially also implicitly encoding spatiotemporal information. This finding as well as mutual information, suggests that **explicit encoding of spatiotemporal information (embedding) may benefit overall performance**.

These findings show that **model performance, especially without diversification, varies significantly across ecological gradients**. This has implications when assessing model performance for downstream tasks such as threshold selection. Evaluation of model performance across strata is both informative of model generalisability and can inform sampling decisions. Such analysis depends on the availability of metadata (e.g. location, date, etc).

Finally, limitations remain inherent to the datasets. Both AnuraSet and WABAD-Europe exhibit imbalanced and incomplete coverage, particularly in temporal strata. Sampling bias and overrepresentation of positive events inflate performance relative to real-world conditions.

## CRediT Authorship Contribution Statement

**Ben McEwen**: conceptualisation, implementation, writing. **Corentin Bernard**: data preparation, implementation and review. **Dan Stowell**: conceptualisation and review.

